# Physico-biochemical and molecular responses of *Acacia auriculiformis* to salinity stress

**DOI:** 10.1101/837351

**Authors:** Deepak B. Gupta, Joydeep Banerjee, Krishnendu Pramanik, Arpita Das, Srikumar Pal

## Abstract

Salinity limits the growth and yield of many crops across the globe and is considered as major threat to agriculture. *Acacia auriculiformis*, an important salt tolerant crop, is growing abundantly in the salt-affected mangrove areas of Sunderban, West Bengal, India. In the present study, we have reported the physiological, molecular and antioxidant response of this crop to a gradient of salt treatments ranging between 0 and 800 mM NaCl. As a stress response, the antioxidant enzymes *viz*. superoxide dismutase (SOD), catalase (CAT) and guaiacol peroxidase (GPX) were highly activated at 200, 400 and 800 mM NaCl respectively. Antioxidant metabolites such as phenols and thiols elevated with increasing salt treatments thus augmenting antioxidant activity with significant positive relationship with phenol content. Similarly, phenylalanine ammonia lyase (PAL) activity was up-regulated in a dose dependent manner with significant relationship with phenol content. This study also reported the phenolic profile for the first time in *A. auriculiformis* with the abundance of flavonoids. In addition, transcriptional up-regulation of Na^+^/H^+^ antiporter gene (*NHX1*) and the development of robust vascular tissues was noticed at 400 mM NaCl stress compared to control, while further stress at 800 mM NaCl induced poor vascular tissue growth but with higher PAL activity and consequent higher phenol content. Based on this observation, a model for salt tolerance mechanism of *A. auriculiformis* has been proposed.

## Introduction

Salinity is known to severely affect the growth and yield of many crops in arid and semi-arid areas of the world covering approximately 20% global cultivated land [1]. Salt stress, in general, causes both hyperionic and hyperosmotic effects in plants, eventually leading to disruption of ion homeostasis, membrane disorganization, excess generation of reactive oxygen species (ROS), which may accelerate stress-induced oxidative damage of biological macromolecules and cellular structure [2]. In addition, salinity with its effect on stomatal closure reduces the carbon dioxide/ oxygen ratio in leaf tissues and limits carbon dioxide assimilation [3]. The major ROS include superoxide radical anion (O_2_^-.^), hydrogen peroxide (H_2_O_2_), hydroxyl radical (OH.) and singlet oxygen (^1^O_2_), which are usually produced at the reaction centers of photosystem I and II (PS I and PS II) in chloroplast [4]. In particular, O_2_^-^ is generated by the non-cyclic electron transport, when stress conditions favours accumulation of nicotinamide adenine dinucleotide phosphate (NADPH) and limit CO_2_ assimilation [5]. Subsequently, O_2_^-^ is converted to hydrogen peroxide (H_2_O_2_) through dismutation reaction catalyzed by SOD and finally to highly oxidizing hydroxyl radical (OH.) through Fenton reaction [6]. Plants have evolved a well defined but complex antioxidant system consisting of enzymatic antioxidants such as SOD, CAT, specific and non-specific peroxidases, glutathione reductase (GR) and metabolites viz. ascorbate, glutathione and phenolic compounds [6,7]. Among these metabolites, phenolic compounds are widespread in plant kingdom and their accumulation is considered as a self-protection way to strengthen ROS scavenging capacity in plants for alleviating salt induced oxidative stress [8,9,10]. Indeed, the enhanced accumulation of phenolic compounds under different environmental stresses triggers upregulation of PAL, the key enzyme of phenol synthesis [11]. Thus, plants under salinity stress can serve as potential source of phenolic compounds with restricted biomass production. Moreover, several plants have evolved diverse anatomical, physiological, biochemical and molecular mechanisms [12], which allow them either to sustain or limit their growth under stressed condition. Halophytes usually overcome salt stress either by removing the cytosolic Na^+^ load by way of transporting it into vacuole via Na^+^/H^+^ antiporter [13] or by accumulating enhanced level of osmo-protectants [14].

*Acacia auriculiformis*, a representative member of Fabaceae, is reputed for its use as fodder and textile industry because of its high tannin content. Moreover, this crop has the potential to find its place in social forestry and cropland agroforestry because of its fast growing nature and wide adaptability to diverse environmental conditions [15]. In West Bengal, India, this crop is grown abundantly in costal salt-affected region of Sunderban and exhibits a high degree of salt tolerance. The underlying salt tolerance mechanism of this crop is reported to be closely associated with ion selectivity, increased accumulation of osmoprotectants, sequesteration of Na in root tissues, anatomical adjustments, Na exclusion and tissue tolerance mechanism [14,16]. However, antioxidant metabolism in response to salinity stress in *A. auriculiformis* is ambiguous and not well documented.

Therefore, the present study was extended to examine the activities of antioxidant enzymes *viz*. SOD, CAT, GPX and PAL in response to a gradient of salinity stress, ranging from 0 to 800 mM NaCl. Moreover, total phenol content with its profile of phenolic compounds and associated antioxidant activity were also explored. Salt induced physiological changes such as cross-linking of cell wall components and vascular tissue development through histological staining have also been noticed. Additionally, gene expression of Na^+^/H^+^-antiportes in *A. auriculiformis* at 400 mM has been reported.

## Materials and Methods

### Salinity treatment and morphological study

The three month-old seedlings of *A. auriculiformis*, collected from salt affected areas of Sunderban, South 24 Parganas, West Bengal, India and grown in sandy-clay soil in pots (25 cm diameter × 20 cm height) in three replicates at the University green house at Mohanpur, Nadia, West Bengal and treated each with 0, 200, 400, 600 and 800 mM Nacl for consecutive 10 days. The growth pattern was analyzed. For biochemical studies, the leaves from each treatment replications were harvested and preserved at (-) 80 °C until further use.

### Assay of antioxidant enzymes

All extraction steps were carried out at 4° C and SOD and CAT were analyzed based on standard protocol [17]. SOD (EC 1.15.1.1) was extracted in 50 mM phosphate buffer (pH 7.8) containing 2% PVP and 0.25 % Triton-X. SOD was assayed in 50 mM phosphate buffer (pH 7.8), 20 µM methionine, 1.12 µM nitroblue tetrazolium (NBT), 1.5 µM EDTA and 75 µM riboflavin using the photochemical NBT method in terms of SOD’s ability to inhibit reduction of NBT to form formazon by superoxide. The photoreduction of NBT was measured at 560 nm. The SOD activity was expressed in units, which were defined as amount of sample required to cause 50% inhibition of photo-reduction of NBT per minute. CAT (EC 1.11.1.6) and GPX (EC 1.11.1.7) were extracted in 100 mM phosphate buffer (pH 7.5) containing 2% PVP and 0.25 % Triton-X. CAT was evaluated spectrophotometrically by determining the consumption of H_2_O_2_ at 240 nm in 100 mM phosphate buffer (pH 7.5) and 200 mM H_2_O_2_. CAT activity was expressed as µmol of H_2_O_2_ destroyed /g/ min. GPX was assayed in 100 mM phosphate buffer (pH 7.5), 30 mM H_2_O_2_ and 2 mM guiaicol. The absorption at 470 nm was recorded and the activity was calculated using the extinction coefficient of 26.6 mM^-1^cm^-1^. GPX activity was expressed as µmol of guiaicol oxidized/ g / min. PAL (EC 4.3.1.5) was extracted in 100 mM phosphate buffer, pH 7.5 containing 2% PVP and 0.25% Triton-X [18]. The enzyme was examined spectrophotometrically by the formation of *t*-cinnamic acid at 270 nm in 1.9 ml of 100 mM Tris-HCl buffer (pH 8.8) and 10 mM phenylalanine. PAL activity is expressed as µmol of *t*-cinnamic acid formed /g / min.

### Analysis of total phenol, phenolic compounds and total thiol

Total phenol was extracted after boiling at 90° C for 2 h with 1.2 M HCl in 50% aqueous methanol [19]. The supernatant obtained after centrifugation was filtered through Whatman 42 filter paper. Total phenol content was determined using Folin-Ciocalteau Reagent (FCR) and expressed as mg gallic acid equivalent per gram dry weight (mg GAE/g DW). For the analysis of phenolic compounds by HPLC, the extract was further filtered through syringe filter (0.22 mm nylon syringe filters; Phenomenex, Torrance, CA, USA). The separation of phenolic compounds was performed in a HPLC (Agilent 1220) coupled with C-18 RP column (125/4 mm with 5 mm particle size), PDA detector (WL 280 nm). The mobile phase consisting of solvent A (1% formic acid in water) and solvent B (1% formic acid in acetonitrile) was used at a flow of 1 ml/ min with the following gradient program: 15% B linear from 0 to 12 min.; 50% B linear from 12 to 35 min.; 85% B linear from 35 to 45 min.; 15% B linear from 45 to 50 min. The column temperature was maintained at 35°C. The sample (20 µL) was injected in duplicate. The chromatographic peaks were identified based on comparison of retention time with those of authentic analytical standards. The quantitative analysis of phenolic compounds was made by chromatographic chemistation software [20] using a linear regression for the relationship of peak area versus concentration of phenolic compound.

Total thiol was extracted in 20 mM EDTA and measured [21] at 412 nm in a reaction mixture consisting of sample extract, 200 mM Tris-HCl buffer (pH 8.O), and 10 mM DTNB following incubation for 15 minutes at room temperature. Total thiol was expressed as µmol of GSH per gram fresh weight of sample (µmol GSH/ g FW).

### Analysis of antioxidant activity

The antioxidant activity of the phenol extract was determined according to earlier reported protocol [22]. In 1,1-diphenyl-2-pycrylhydrazyl (DPPH) assay, the changes in absorbance was measured at 517 nm in reaction mixture containing 150 µl of phenolic extract and 2850 µl of 0.004% DPPH while that in Ferric Reducing Antioxidant Power (FRAP) assay was recorded at 593 nm in reaction mixture consisting of 150 µl neutralized sample extract and 2850 µl FRAP reagent. The antioxidant activity was expressed as mg trolox equivalent per gram dry weight (mg TE/g DW).

### Analysis of NHX1 gene expression in A. auriculiformis

Transcript level expression study of Na^+^/ H^+^ antiporter (*NHX1*) in *A. auriculiformis* was carried out considering the forward primer (*NHX1*F; 5′-TTTGGCAGGCACTCAACAGA-3′) and reverse primer (*NHX1*R; 5′-TAAGGAGTCCTGCCCCTCTC-3′) in semi-quantitative reverse transcriptase PCR using total RNA isolated from leaf tissue of control and 400 mM NaCl treated plants at same developmental stages. Actin gene specific primer (*Act*F; 5′-GTGCCCATTTACGAAGGATA-3′ and *Act*R; 5′-GAAGACTCCATGCCGATCAT-3′) were used as endogenous loading control.

### Pholoroglucinol Staining

The transverse section of freshly harvested phyllode, obtained from previously treated *A. auriculiformis* with 0, 400 and 800 mM Nacl was subjected to phloroglucinol staining following the protocol reported earlier [23].

### Statistical analysis

Experimental results were expressed as means ± SD considering three replicates for all the measurements. The data were analyzed by an analysis of variance (p < 0.05) and significant differences among the treatments were tested by Duncan’s multiple range test at 5% level using Statistical Tool for Agricultural Research (STAR) software package version 2.0.1 of International Rice Research Institute (IRRI). Correlation coefficients (r) were also calculated between antioxidant activity and phenol content.

## Results

### Effect of salt stress on SOD, CAT, GPX and PAL

Salt treatments significantly enhanced SOD activity (Fig. 1A) over control at 200 and 400 mM NaCl, but the stimulatory effect was more pronounced at 400 mM. On the contrary, at 600 and 800 mM salt treatment, SOD activity declined significantly below control. Hence, the SOD activity might be either up-regulated or down-regulated depending on the severity of stress. The result however, suggested that relatively higher H_2_O_2_ generation was imposed at lower (200 and 400 mM) than higher (600 and 800 mM) salt concentration. The CAT activity (Fig. 1B), on the other hand, increased significantly over control with all the treatments except 400 mM. It was further noticed that PAL (Fig. 1C) activity and GPX activity (Fig. 2A), the key enzymes of phenol synthesis and oxidation respectively, controlling the phenol accumulation in plants, increased significantly over control in response to increasing salt treatments.

**Fig. 1.**
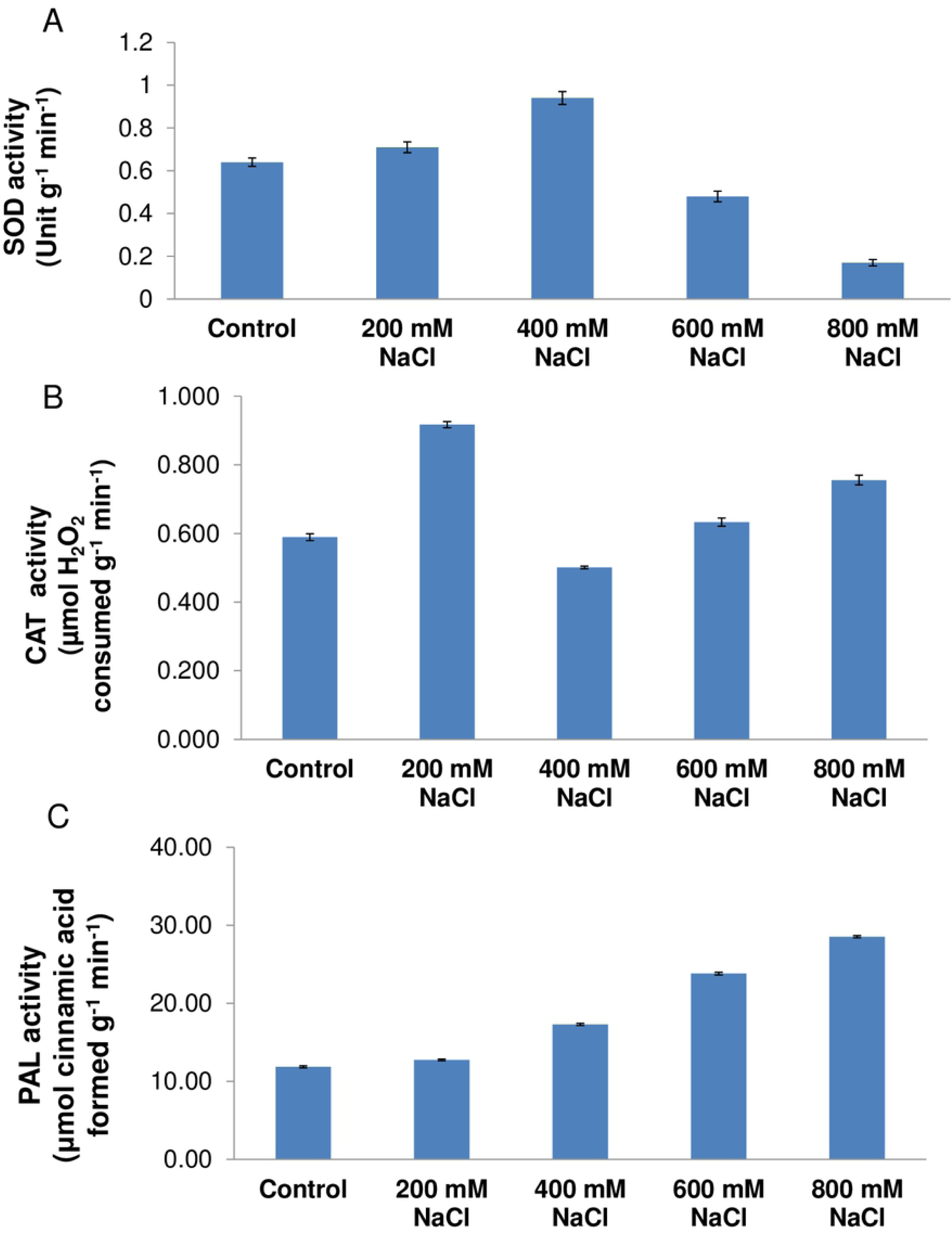
Effect of salt stress on enzymatic antioxidants in *A. auriculiformis*. (A) Superoxide dismutase (SOD) activity in leaf (phyllode) tissue of untreated (Control) and indicated amount of NaCl stressed seedlings. (B) Catalase (CAT) activity in leaf (phyllode) tissue of untreated (Control) and indicated amount of NaCl stressed seedlings. (C) Phenylalanine ammonia-lyase (PAL) activity in the leaf (phyllode) tissue of untreated (Control) and indicated amount of NaCl stressed seedlings. Data represent mean± SD having n = 3.

**Fig. 2.**
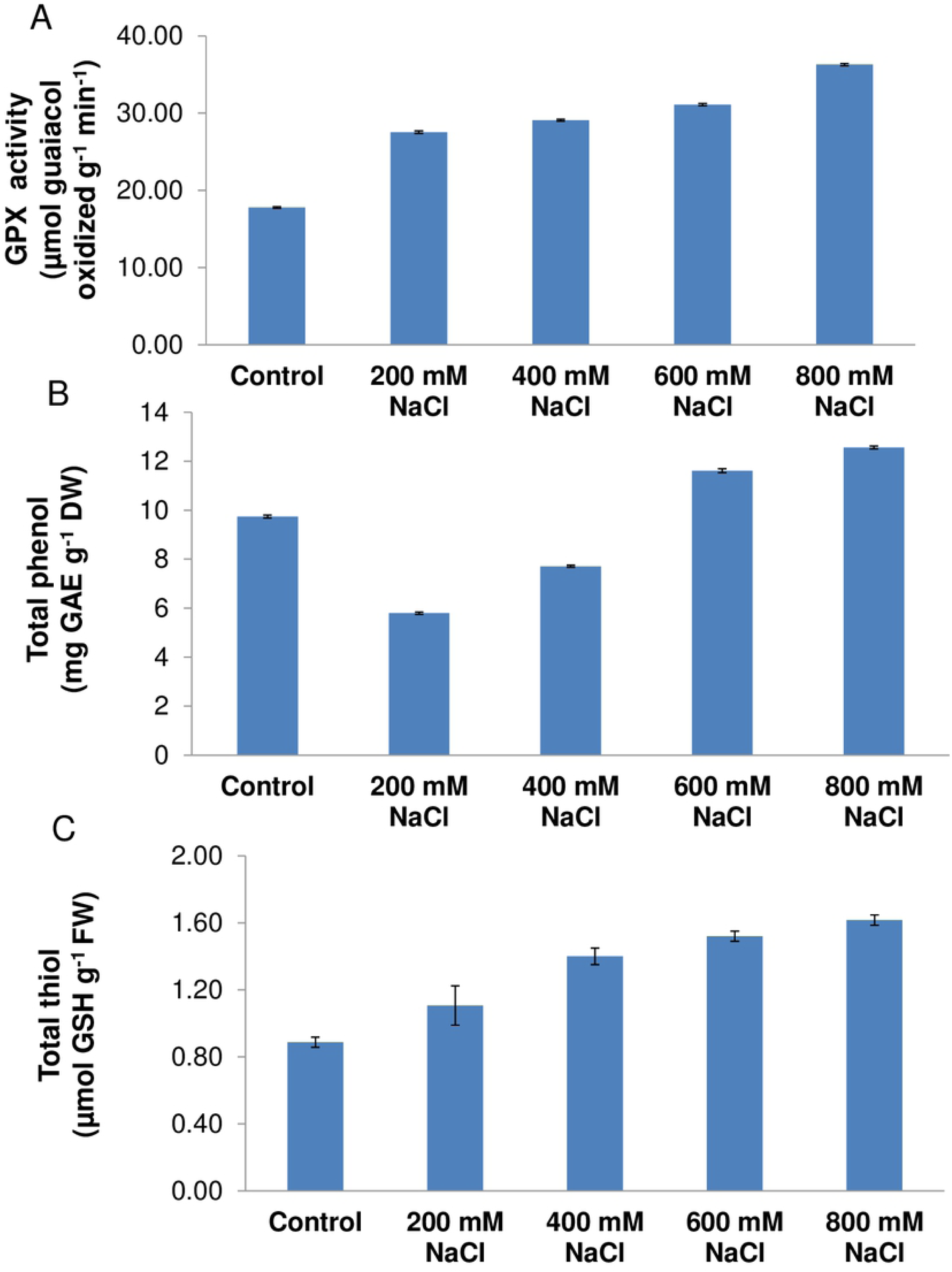
Effect of salt stress on non-enzymatic antioxidants, PAL activity and total antioxidant activity in *A. auriculiformis*. **(A)** Guaiacol peroxidase (GPX) activity in leaf (phyllode) tissue of untreated (Control) and indicated amount of NaCl stressed seedlings.Total phenol (B) and total thiol (C) content in the leaf (phyllode) tissue of untreated (Control) and indicated amount of NaCl stressed seedlings.

### Effect of salt treatment on content of phenolic compounds and reduced glutathione

Total phenol content (Fig. 2B) increased significantly over control with increasing level of salt treatments. Similar to phenol, total thiol content increased significantly with increasing salinity (Fig. 2C). In the present study an attempt has been made for analysis of 16 phenolic compounds (Fig. 3) consisting of 4 hydroxybenzoic acids *viz*. gallic (Gal), protocatechuic (Pro), p-hydroxybenzoic (Pba) and vanillic acid (Van), 5 hydroxycinnamic acids *viz*. p-comaric (Pco), caffeic (Caf), ferulic (Fer), cinnamic (Cin) and chlorogenic acid (Chl) and 7 flavonoids *viz*. myricetin (Myr), apigenin (Api), quercetin (Qur), catechin (Cat), luteolin (Lut) Rutin (Rut) and kaemferol (Kam). However, our study documented the detection of 15 phenolic compounds in *A. acquiformis* (Table 1). The flavonoids formed the major class of phenolic compounds followed by hydroxybenzoic acid representing 66.55 and 27.34% respectively of total phenolic compounds. Among the phenolic compounds, catechin was observed as most abundant forming 51.17% of total phenolic compounds and its content increased significantly in a dose dependent manner. Gallic acid ranked distinct second representing nearly 16% of total phenolic compounds. Salt treatment with 200 and 400mM NaCl did not produce any significant differences in gallic acid content with the control. However, gallic acid content at 600 and 800 mM has found to differ significantly not only between them but also from other treatments. Protocatechuic acid with its share of 6.28% formed the third major phenolic compound and increased significantly at higher treatments (600 and 800 mM). The next abundant compound was chlorogenic acid, an ester of caffeic and quinic acid and increased significantly with increasing salt treatments. The phenolic compound detected as minor included p-hydroxybenzoic, caffeic, p-coumaric ferulic and cinnamic acid. The mean of vanillic acid and other flavonoids together formed 19.12% of total phenolic compound.

**Table. 1:** Mean value of 15 phenolic compounds detected under various salt concentration in *A. acquiformis*

| Sl No. | Parameter | Treatment |  |  |  |  |
| --- | --- | --- | --- | --- | --- | --- |
|  |  | A0 | A200 | A400 | A600 | A800 |
| 1. Hydroxyl Benzoic Acid (µg/g) DW |  |  |  |  |  |  |
| a. | Gallic Acid | 558.46 c | 556.64 c | 560.14 c | 578.44 b | 606.88 a |
| b. | Proto catechuic | 164.14 d | 218.76 c | 222.78 c | 238.72 b | 284.52 a |
| c. | P.Oh. Benzoic | 36.13 d | 39.35 d | 44.91 c | 56.78 b | 76.12 a |
| d. | Vanillic Acid | 88.23 d | 102.55 c | 154.35 b | 156.98 b | 171.35 a |
|  | Total | 846.96 e | 917.3 d | 982.18 c | 1030.92 b | 1138.87 a |
| 2. Hydroxy Cinnamic Acid (µg/g) DW |  |  |  |  |  |  |
| a. | Caffeic Acid | 48.54 d | 52.76 d | 60.58 c | 72.67 b | 81.37 a |
| b. | Cinnamic Acid | 2.17 c | 2.76 b | 3.69 a | 3.57 a | 2.04 d |
| c. | P. Coumaric | 2.37 e | 3.14 d | 6.68 a | 5.07 b | 4.55 c |
| d. | Cholorogenic Acid | 106.34 d | 124.67 c | 154.32 b | 161.76 b | 174.65 a |
| e. | Ferulic Acid | 6.98 a | 5.05 c | 6.21 b | 4.66 d | 3.38 e |
|  | Total | 166.4 e | 188.38 d | 231.48 c | 247.73 b | 265.99 a |
| 3. Flavonoids (µg/g) DW |  |  |  |  |  |  |
| a. | Myricetin | 49.77 d | 88.73 c | 118.44 b | 132.96 a | 138.34 a |
| b. | Luteolin | 82.08 c | 77.98 c | 122.68 b | 116.96 b | 132.12 a |
| c. | Kaemferol | 78.22 d | 108.55 c | 117.71 b | 114.78 b | 126.29 a |
| d. | Rutin | Not detected | Not detected | Not detected | Not detected | Not detected |
| e. | Quercetin | 74.56 e | 99.54 d | 132.11 c | 146.76 b | 156.67 a |
| f. | Catechin | 1696.12 e | 1743.66 d | 1858.62 c | 1906.77 b | 1996.77 a |
| g. | Apigenin | 74.58 e | 90.43 d | 108.45 c | 128.43 b | 148.32 a |
|  | Total | 2055.33 | 2208.89c | 2458.01b | 2546.66b | 2698.51a |
|  | Grand Total | 3068.69e | 3314.57d | 36791.67c | 3825.31b | 4103.37a |

Different letters indicate significant difference at $P < 0.05$ (or means within columns separated by Duncan's Multiple Range Test $P = 0.05$ [44].

**Fig. 3.**
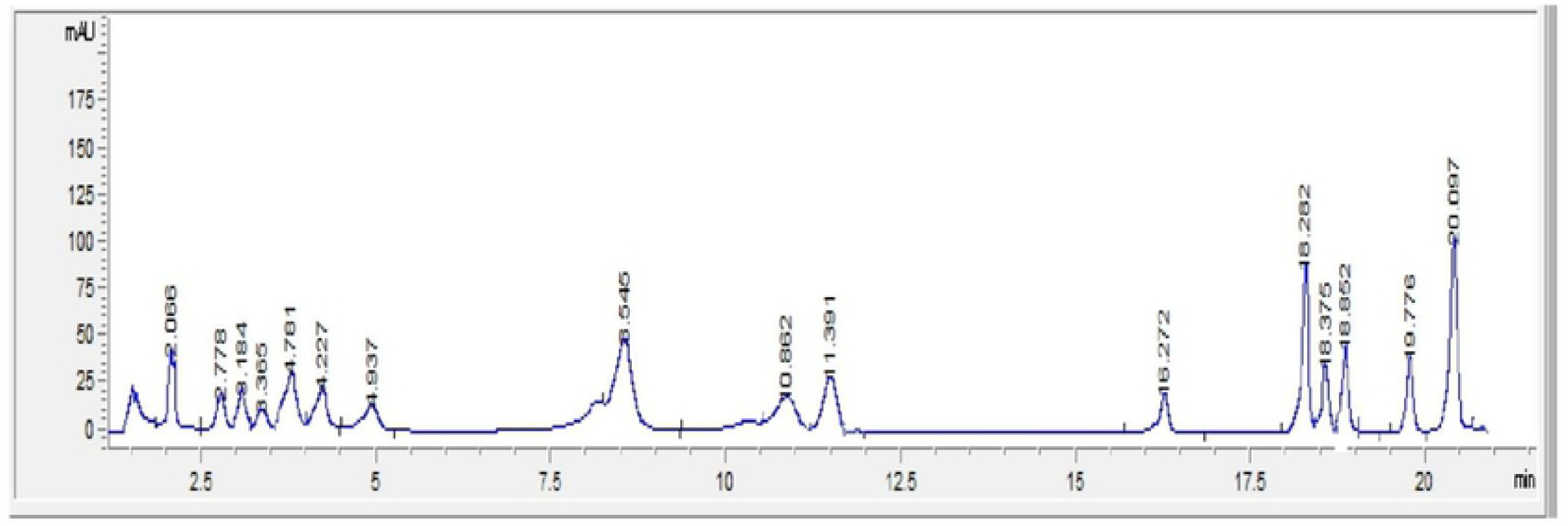
Retention time of 16 phenolic compounds (Retention time gallic acid:2.063, protocatechuic acid:2.774, chlorogenic acid:3.174, catechin:3.357, p-hydroxybenzoic acid:4.217, caffeic acid:4.75, vanillic acid:4.929, p-comaric acid:8.512, ferulic acid:10.837, rutin:11.291, myricetin:16.251, luteolin:18.266, quercetin: 18.376 cinnamic acid:18.836, apigenin:19.761, kaemferol:20.059 min) detected in *Acacia auriculiformis*.

### Effect of salt treatment on antioxidant activity

The antioxidant activity, measured under DPPH and FRAP assay (Fig. 4), was significantly increased over control with increasing salt treatments. However, the antioxidant activity measured under DPPH assay produced a greater absolute value than that obtained from FRAP assay. This difference indicated that phenolic compounds occurring in *A. auriculiformis* scavenged ROS chiefly by hydrogen atom transfer (HAT) rather than single electron transfer (SET) mechanism. The antioxidant activity showed a significant and positive relationship with phenol in DPPH (r= 0.73) and in FRAP assay (r= 0.71).

**Fig. 4.**
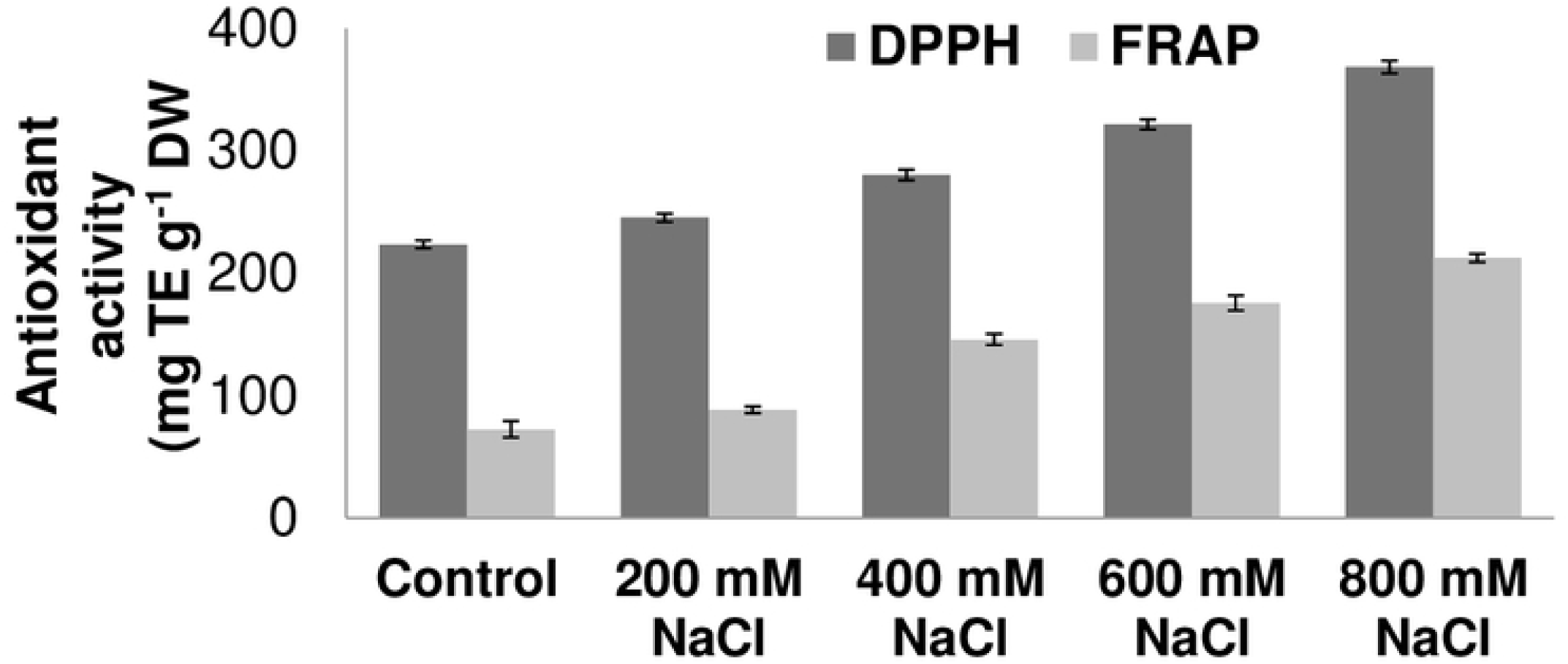
Total antioxidant properties under DPPH and FRAP assay in the leaf (phyllode) tissue of untreated (Control) and indicated amount of NaCl stressed seedlings of *A. auriculiformis*. Data represent mean± SD having n = 3.

### Salt stress induced physiological, anatomical and gene expressional changes

Application of salt treatment at varying concentration resulted in differential physiological response (Fig. 5A). In untreated control, more number of leaves occasionally in one branch was noticed after 10 days, while the number of leaves declined with increasing salt treatment except at 800 mM NaCl where, appearance of new leaves was noticed. Moreover, some curled leaves were also detected at 600 mM NaCl, while 800 mM salt treatment resulted in almost curved leaves. Morphological study clearly demonstrated that dry matter accumulation in seedling of *A. auriculiformis* decreased with increasing salt treatment.

**Fig. 5.**
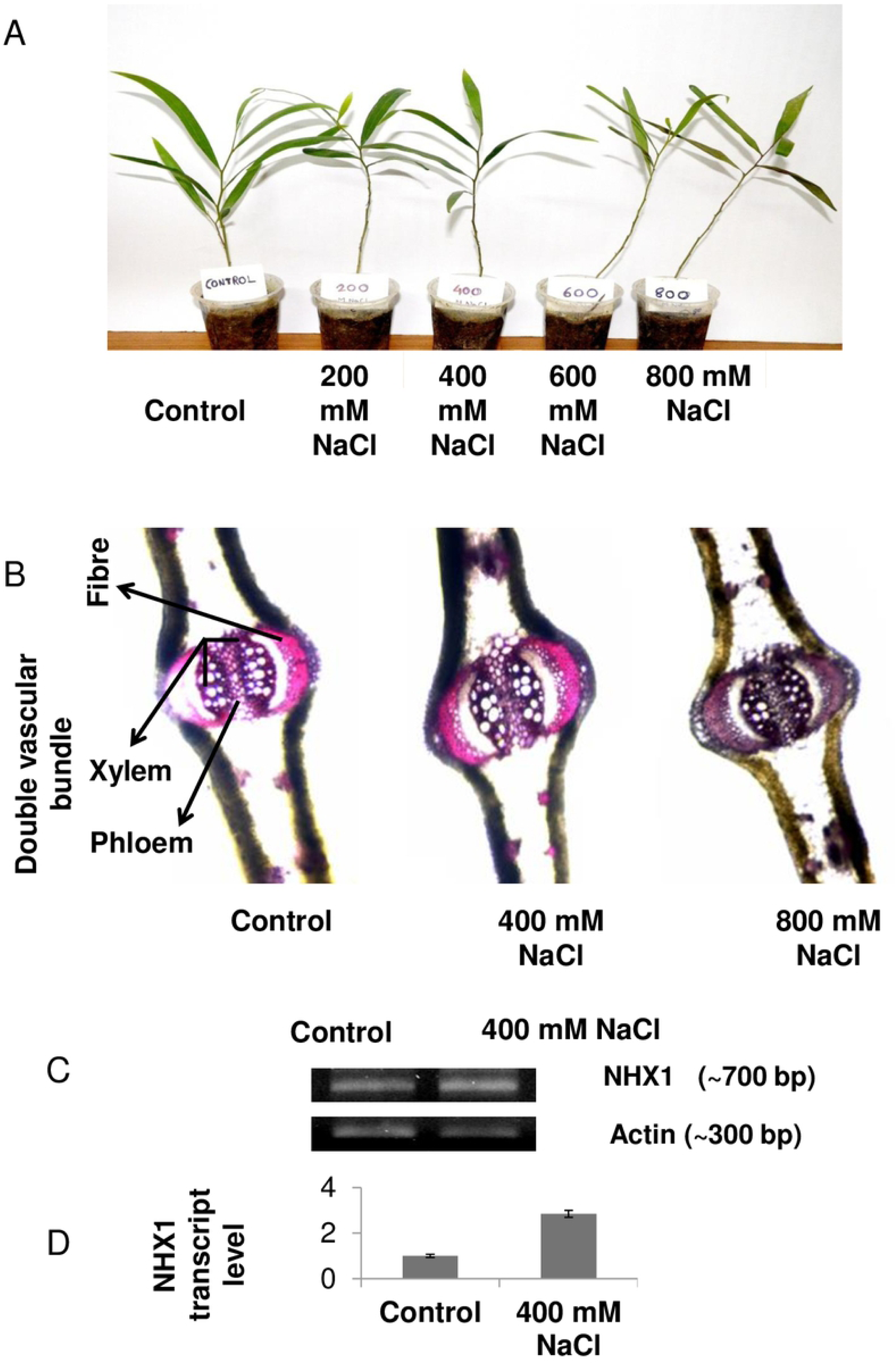
Morphological, anatomical and molecular changes in *A. auriculiformis* upon salt stress. (A) Morphological study on *A. auriculiformis* seedlings after indicated amount of salt stress for 10 days along with deionized water treated control seedling. (B) Phloroglucinol staining of transverse section of *A. auriculiformis* phyllode grown in control and indicated amount of salt stressed condition for 10 days. Bar indicates 200 μm. (C) Transcript expression of *NHX1* in the leaf tissue of control and 400 mM NaCl treated *A. auriculiformis*. Semi-quantitative RT-PCR of control and 400 mM NaCl treated *A. auriculiformis* seedlings using *Actin* as internal control. (D) The relative expression of *NHX1* gene (calculated using ImajeJ software analysis taking *Actin* gene expression as loading control) in control and 400 mM NaCl treated *A. auriculiformis* seedlings. Values are means ±SD with n = 3.

Differential salt stress depicted anatomical changes in *A. auriculiformis* (Fig. 5B). Pholoroglucinol staining documented significant changes in vascular tissues across different treatments. Pholoroglucinol staining established more cross-linking of cell wall components with 400 mM NaCl but less with 800 mM NaCl treated phyllode over control. In addition, 400 mM NaCl stressed phyllode exhibited better vascular tissue development as opposed to 800 mM NaCl stressed condition and even better over control. Additionally, the phloem tissue development in the phallode was found to increase in the order of 800 mM < control < 400 mM NaCl treated seedlings. As 400 mM NaCl treatment depicted more cross-linking in seedling compared to control as well as 800 mM NaCl treatment, the transcript expression of a Na^+^/H^+^ antiporters gene (*NHX1*) was analyzed in 400 mM NaCl treated seedlings. Relative transcript abundance of *NHX1* gene was found to be up regulated at 400 mM NaCl stress compared to untreated control plants (Fig. 5C). Quantification of relative expression of Na^+^/H^+^ antiporters (relative to Actin) conducted through imajej software analysis revealed that upon 400 mM NaCl stress the gene was 2.84 fold up-regulated compared to control plants (Fig. 5D).

## Discussion

All plants under suboptimal environmental condition, facing abiotic stress, experience an imbalance between the energy received and the ability to process it. ROS formation, photo-inhibition and growth reduction are the major consequences of such energy imbalances [24,25]. In the present study, decline in number of leaves observed with increasing salinity suggested that the growth reduction appears to be related to impairment of either non-cyclic photosynthetic electron transport or CO_2_ assimilation. Earlier study reported that the growth related parameters including stomatal conductance, photosynthetic pigments, protein and carbohydrate content in *A. auriculiformis* were adversely affected by salinity in a time and dose dependent manner [14]. Although, measurement of MDA content serves as an indicator of ROS mediated oxidative injury in plant cells, the differential stimulation of SOD and CAT activity to the severity of salt stress provides an indirect evidence of generation of ROS. In the present study comparison of SOD and CAT activity across different salt treatments, provided a clue to greater accumulation of H_2_O_2_ at 400 mM, which was also evidenced by higher SOD and moderate CAT activity. Similar activation of SOD was noticed at 400 mM in a halophyte, *Bruguiera pariflora*, growing abundantly in salt affected areas of Sunderban, West Bengal, India [26]. On the contrary, the moderate SOD and higher CAT activity at 800 mM resulted in lower concentration of H_2_O_2_. Relatively lower SOD activity at higher salt stress was similar to the previous report in shoot tissues of *Brassica napus* L [27]. In rice, it was observed that SOD activity was decreased at higher salt concentration in sensitive rice genotypes in contrast to stimulatory effect at lower concentration in tolerant genotypes [28]. Similar increase in CAT activity was also reported for salt-tolerant cotton [29].

In the present study, the enhanced GPX activity with increasing salinity was corroborated with previous reports in rice [30] and other plants [31,32]. The enhanced GPX activity, which uses H_2_O_2_ as a substrate for oxidation of phenolic compounds further reduced the H_2_O_2_ concentration at 800 mM. Thus, it appeared reasonable to assume that the most toxic ROS, the OH^-^, is formed via Fenton-like reaction, in greater abundance at 400 than 800 mM NaCl. Consequently, ROS generation in *A. auriculiformis* in response to salt treatment revealed higher with 400 mM than 800 mM NaCl. Moreover, higher thiol content at 800 mM than 400 mM salt stress further reduced the ROS load at 800 mM as these compounds were reported to scavenge ROS *in vivo*. Moreover, total thiol primarily comprises of reduced GSH, a major component of thiol compounds act as a substrate for glutathione peroxidase and thus reduces the H_2_O_2_ load in cell. In general, environmental stresses and elicitor are known to modulate the phenolic compounds in plants [33,34] and changes in these compounds are critically associated with salt sensitivity of plant as well as severity of salt stress [35,36]. They traditionally appear in leaves with obvious oxidative stress. Therefore, leaves of *A. auriculiformis* upon exposure to 400 mM NaCl stress should accumulate greater phenol. By contrast, leaf treated with 800 mM NaCl and with consequent lower OH radical concentration, displayed significantly higher PAL activity, phenol concentration and in vitro antioxidant activity. In corroboration with the previous report, it disclosed that inhibition of CO_2_ assimilation at 800 mM salt stress resulted greater NADPH accumulation through linear photosynthetic electron transport, which is reversed by the consumption of NADPH for phenol synthesis [14]. Similar to our finding, salinity induced enhanced accumulation of phenol has been observed in radish [37], pepper [38], honeysuckle [39] and *Cakile maritima* [40]. Moreover, higher *in vitro* antioxidant activity associated with higher phenol concentration, though compared well with the earlier reports [40] had hardly any chance to scavenge ROS *in vivo* in absence of oxidative stress [36,41]. The higher accumulation of phenolic compounds with increasing salinity, flavonoids in particular, act as energy escape valve rather than antioxidant function, to dissipate excess excitation energy resulting from salinity stress [42]. With increasing salinity, the higher accumulation of phenolic compounds, flavonoids in particular, with their role in energy dissipation and carbon diversion, act as energy escape valve rather than antioxidant function [42].

Overproduction of H_2_O_2_ at 400 mM generated not only ample OH radical concentration but also lignin precursor through GPX reaction. All these events, in turn, facilitated ROS mediated cross-linking in cell wall polymeric components and support our observation of phloroglucinol staining. In corroboration to our present findings, earlier study documented that strong vascular tissue development is helpful for the seedlings to withstand salt stress in a better manner [43]. The antioxidant response at 400 mM NaCl thus improved the cell wall integrity and membrane permeability that might lead to greater accumulation of Na^+^ in cytosol. However, the over expression of *NHX1* gene at 400 mM was triggered to reduce cytosolic Na concentration by allowing its transportation into vacuole. Similar to our finding, salinity induced up-regulation of transcription of *NHX1* gene in several tree species including *Populus euphortica* and *Aluropus littoralis* has been reported [44]. In contrast, antioxidant response with higher phenol accumulation with its role in photoprotection provided a back-up defense at 800 mM.

Based on our finding, a plausible model for salt tolerance in *A. auriculiformis* has been proposed (Fig. 6). In addition to adverse effect on the growth depending on the severity of stress, central to this model was the activation of SOD and PAL. At 400 mM, higher activation of SOD accompanied by moderate CAT activity was the probable reason for enhanced production of H_2_O_2_ followed by higher OH^-.^ load in plant cell. H_2_O_2_ with its involvement as a substrate in GPX reaction act as lignin precursor and facilitate cross-linking of cell wall materials. Moreover, over expression of *NHX1* gene at 400 mM reduced hyper-ionic Na^+^ stress via its transport to vacuole. On the other hand, at 800 mM the activation of PAL accompanied by greater phenol accumulation resulted in lower ROS due to involvement of NADPH for phenol synthesis and its photo protective role. However, higher phenol accumulation resulting from greater stimulation of PAL at 800 mM salt stress provided a mechanism to reduce the energy imbalance between that received and processed by plant by dissipating it as either as heat or fluorescence or both. Present study judiciously unfolds the physico-biochemical and molecular mechanism behind the salt stress response in *A. auriculiformis* and affirms the significance of this tree species for uplifting livelihood and sustenance of salinity affected coastal areas.

**Fig. 6.**
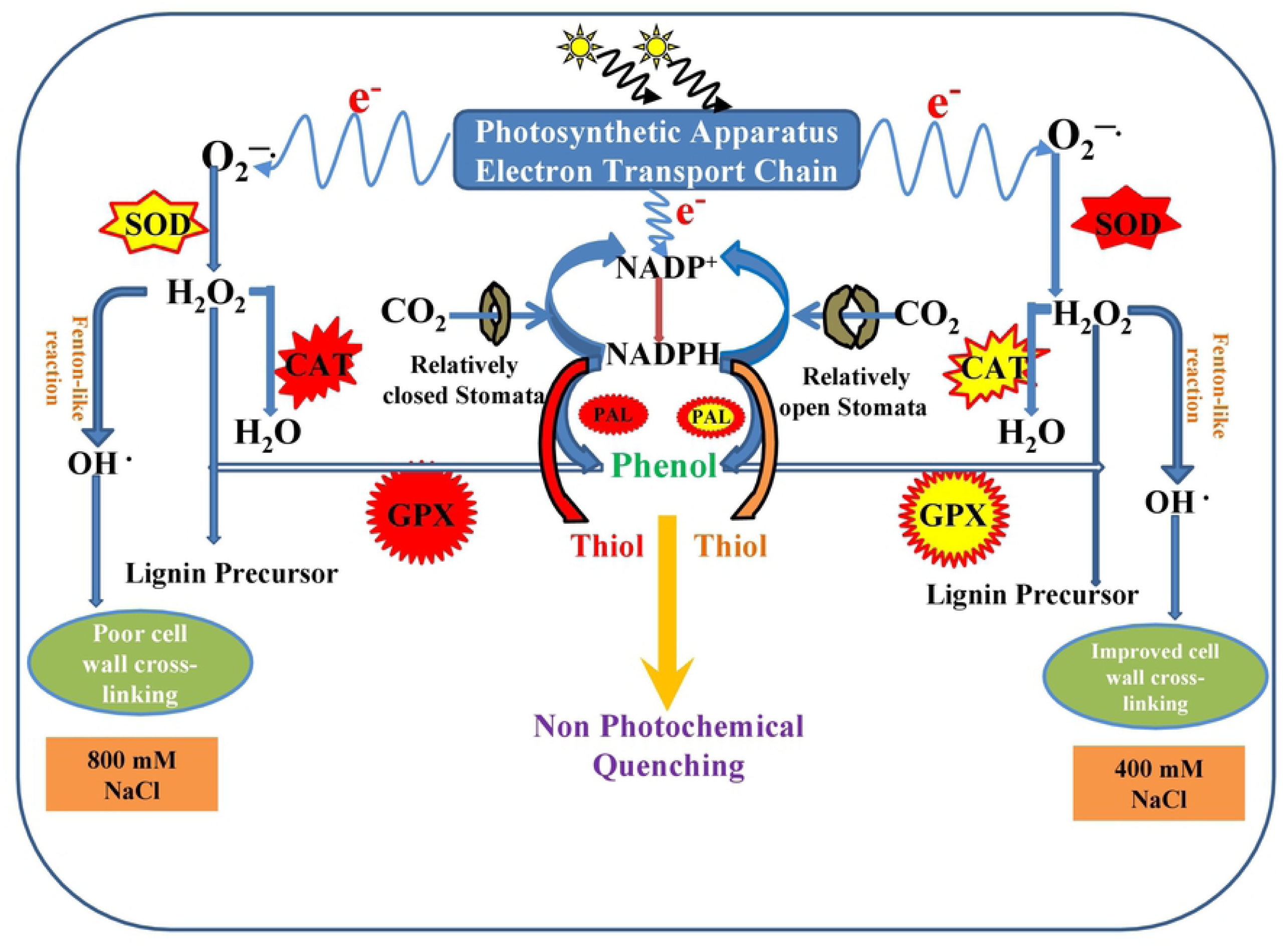
Biochemical cross-talk in *A. auriculiformis* under salt stress. Proposed model depicts the involvement of various enzymatic and non-enzymatic antioxidants under control, 400 mM NaCl stress and 800 mM NaCl stress for the cross-linking of cell wall components. Red and blue arrow indicates enhanced and retarded activity, respectively.

## Acknowledgement

Authors would like to acknowledge Bidhan Chandra Krishi Viswavidyalaya (BCKV) for providing basic support to carry out the research.

## Authors’ Contributions

SP and JB conceptualized the study, DG, KP implemented and executed the experiment, DG, JB and KP collected the data, AD and SP carried out data analysis and interpretation and SP, JB, KP and AD composed the manuscript.

